# Unified integration of spatial transcriptomics across platforms

**DOI:** 10.1101/2025.03.31.646238

**Authors:** Ellie Haber, Ajinkya Deshpande, Jian Ma, Spencer Krieger

## Abstract

Spatial transcriptomics (ST) has transformed our understanding of tissue architecture and cellular interactions, but integrating ST data across platforms remains challenging due to differences in gene panels, data sparsity, and technical variability. Here, we introduce Lloki, a novel framework for integrating imaging-based ST data from diverse platforms without requiring shared gene panels. Lloki addresses ST integration through two key alignment tasks: feature alignment across technologies and batch alignment across datasets. Optimal transport-guided feature propagation adjusts data sparsity to match scRNA-seq references through graph-based imputation, enabling single-cell foundation models such as scGPT to generate unified features. Batch alignment then refines scGPT-transformed embeddings, mitigating batch effects while preserving biological variability. Evaluations on mouse brain samples from five different technologies demonstrate that Lloki outperforms existing methods and is effective for cross-technology spatial gene program identification and tissue slice alignment. Applying Lloki to five ovarian cancer datasets, we identify an integrated gene program indicative of tumor-infiltrating T cells across gene panels. Together, Lloki provides a robust foundation for cross-platform ST studies, with the potential to scale to large atlas datasets, enabling deeper insights into cellular organization and tissue environments.

## Introduction

In single-cell RNA sequencing (scRNA-seq) analysis, dataset integration is crucial for enabling robust comparisons across studies and conditions, with its importance growing alongside large public data repositories such as the Single Cell Portal [1], CellxGene [2], and the Human Cell Atlas [3]. Similarly, integrating spatial transcriptomics (ST) datasets, which contain both spatial coordinates and gene expression, enables comparative analysis across samples, technologies, and conditions, revealing cellular spatial organization and dynamics across diverse contexts in health and disease [4]. Although batch integration has been extensively studied in scRNA-seq [5], ST presents unique challenges. A key difficulty is variation in gene panels across samples, even within the same technology. Moreover, ST technologies differ significantly in sensitivity to specific genes [6], and varying sparsity levels of the data further complicate integration. The goal of ST batch integration is to learn a spatially aware embedding function that maps gene expression profiles into a shared feature space, ensuring biologically similar cells remain close, regardless of technology or gene panel.

Existing ST integration methods, such as STAligner [7], SPIRAL [8], DeepST [9], and PRECAST [10], predominantly align tissue slices based on shared genes, limiting their ability to integrate datasets from platforms with differing gene panels. This challenge is further compounded when integrating multiple datasets, as the intersection of gene sets across technologies becomes increasingly small. For example, our results show that datasets from five different technologies share only 19 genes, despite the smallest gene panel containing over 250 genes.

Recently, single-cell foundation models (scFMs) such as UCE [11], Geneformer [12], scGPT [13], and others [14–16] have emerged, leveraging large-scale scRNA-seq data to learn robust, generalizable cell representations. While promising for scRNA-seq batch integration, attempts to train foundation models specifically on ST data remain limited [17, 18] due to challenges such as limited data, non-overlapping gene panels, and pronounced batch effects. Despite this, scFMs remain attractive for ST integration due to their extensive pretraining on full transcriptomes, offering a scalable solution to variability in gene panels across ST technologies. However, a key limitation is that scFMs’ cell representations are learned almost exclusively from scRNA-seq data, and the differences in sparsity and gene capture efficiency between scRNA-seq and ST data further complicate their direct application to ST integration.

Here, we introduce Lloki, a novel framework for scalable ST integration across diverse technologies without requiring shared gene panels. The framework consists of two key components: (1) Lloki-FP, which leverages optimal transport and feature propagation to transform ST gene expression profiles, aligning their sparsity with scRNA-seq to optimize scGPT embeddings; and (2) Lloki-CAE, a conditional autoencoder that integrates embeddings across ST technologies using a novel loss function balancing batch integration with the preservation of biological information from Lloki-FP embeddings. This unique combination ensures both feature and batch alignment, enabling robust ST data integration while preserving biological specificity and local spatial interactions. We reveal the fragility of relying on shared gene panels for integration and demonstrate that Lloki significantly outperforms state-of-the-art methods across standard batch integration metrics. Additionally, we show that Lloki’s embeddings facilitate important downstream tasks such as physical slice alignment and cross-gene-panel spatially variable gene program identification, enabling cross-technology analysis of tissue structure and organization across diverse contexts.

## Results

### Overview of Lloki

In Lloki, we introduce a novel framework for ST integration by decomposing the task into two alignment problems: (1) feature alignment across gene panels, and (2) batch alignment across technologies.

For feature alignment with Lloki-FP, we leverage scGPT [13] to embed data into a shared feature space, independent of gene panel differences. However, scGPT – and most existing scFMs – only process non-zero expression genes, making high sparsity problematic, as ST data falls outside the model’s scRNA-seq pretraining distribution. Lloki-FP adjusts ST data sparsity by incorporating spatial information to align the data with an scRNA-seq reference, ensuring compatibility with pretrained scFMs. To achieve this, we first calculate a target sparsity for each cell using optimal transport, aligning ST data sparsity with the scRNA-seq reference. We then employ a novel feature propagation method that integrates gene similarity and spatial proximity to impute missing features while preserving biological variation. The sparsity-aligned ST gene expression is then processed by scGPT, generating feature-aligned embeddings for each cell.

For batch alignment with Lloki-CAE, we use a conditional autoencoder to integrate data across ST technologies, using Lloki-FP embeddings as input features for each cell. Our integration strategy combines three loss functions: (1) reconstruction loss to preserve accurate data representations, (2) triplet loss to enhance clustering and mitigate batch effects, and (3) a new biological conservation loss that maintains the local neighborhood structure from the pre-integration embedding space, ensuring that cell-type relationships established by Lloki-FP are preserved during batch correction. Together, these loss functions enable robust integration while preserving cell type clusters across diverse ST platforms.

Lloki introduces several key innovations that distinguish it from existing ST integration methods. Unlike previous approaches that rely on shared gene panels, Lloki-FP accommodates any gene panel, ensuring all available data contributes to biologically meaningful embeddings. Lloki also uniquely in-corporates spatial information to address differences in technical dropout across technologies without over-smoothing, preserving cell-type specificity rather than collapsing cells into region-specific clusters. Additionally, the three-part loss function in Lloki-CAE maintains biologically meaningful heterogeneity while enabling robust batch integration. Finally, Lloki is highly scalable and parallelizable – after initial training, each ST slice is processed individually, and Lloki embeddings can be computed in under 5 minutes using less than 1 GB of GPU memory.

### Lloki enables cross-technology batch integration

We evaluated Lloki’s performance on cross-technology batch integration using one slice from each of five imaging-based ST datasets of coronal mouse brain sections: MERFISH [19] (1,122 genes), MER-SCOPE [20] (550 genes), STARmap [21] (1,022 genes), CosMx [22, 23] (960 genes), and Xenium [24] (248 genes). Despite the smallest panel containing 248 genes, these platforms share only 19 genes, making integration particularly challenging for existing alignment methods. To benchmark Lloki, we compared it against six baseline methods: (1) PCA, (2) Harmony [25] (for scRNA-seq integration), (3) Seurat [26] (for scRNA-seq integration), (4) scVI [27] (for scRNA-seq integration), (5) STAligner [7] (for ST integration), and (6) the raw output of Lloki-FP (prior to batch alignment via Lloki-CAE).

We visualized the embedding space from each method using UMAP (**Fig**. 2a), coloring cells either by batch or cell type. For cell type visualizations, annotations were consolidated into broader labels for consistency across datasets (**Table** S2); for the Xenium dataset, which lacked annotations, cell types were inferred by marker gene analysis (**Supplemental Information**).

**Figure 1:**
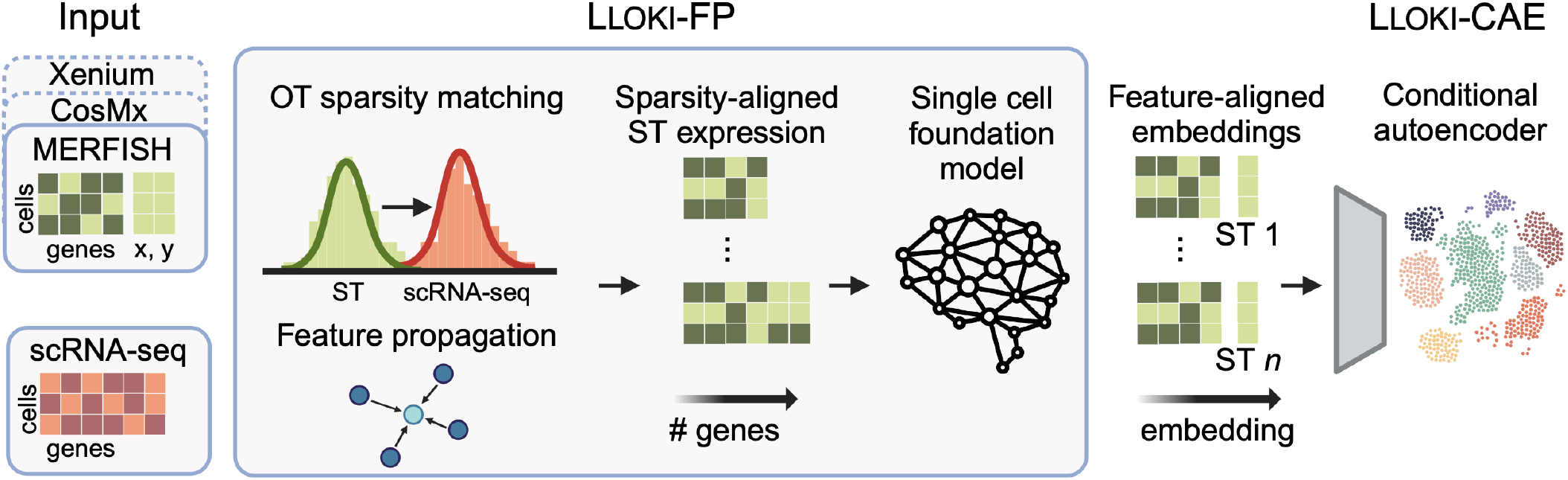
Overview of Lloki. Lloki performs a two-stage alignment for integrating ST data: first, it unifies the feature space across ST samples, regardless of gene panel, then it removes technology-specific batch effects. Lloki-FP applies optimal transport-guided feature propagation to impute missing gene expression values using spatially informed cell graphs, mitigating data sparsity and gene sensitivity differences. A single-cell foundation model then embeds the data into a unified feature space. Lloki-CAE further integrates these embeddings across batches using a conditional autoencoder, removing technology-specific effects while preserving biological structure. The resulting embeddings minimize batch effects and support downstream tasks such as cross-technology slice alignment and spatial gene program identification.

**Figure 2:**
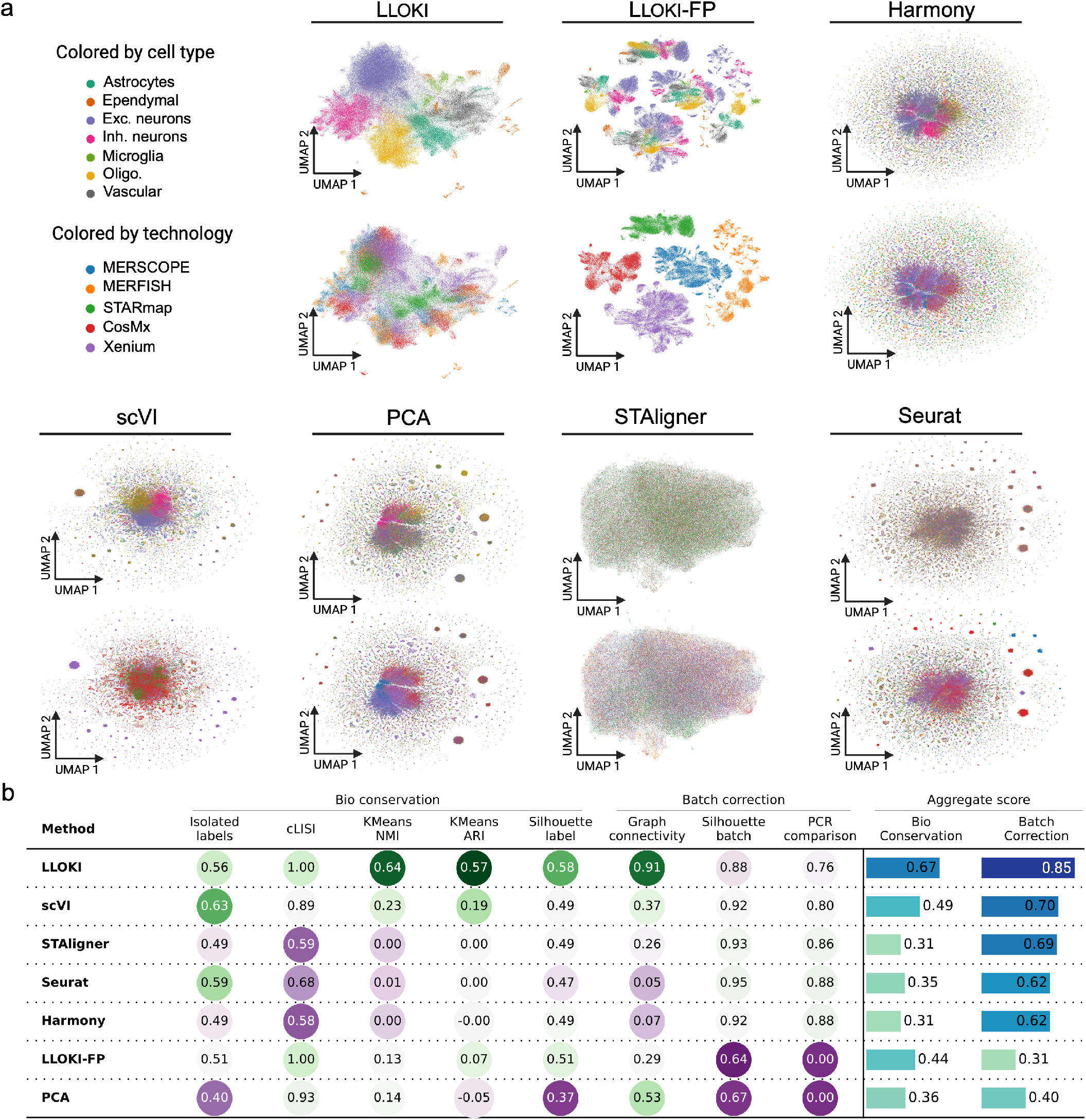
Comparative analysis of batch integration using Lloki versus six baseline methods across slices from five spatial transcriptomics technologies. **a.**UMAP visualizations of integrated data, colored by cell type (top) and by technology (bottom), showing the degree of batch separation and preservation of biological variation. **b**. Quantitative performance evaluation of all methods across eight metrics, measuring both biological variation preservation and batch correction across technologies.

For quantitative evaluation, we employed eight metrics from scib-metrics [28] (**Methods**). Five metrics assessed the preservation of biological variation based on cell type separation using the unified high-level labels for each dataset, and three metrics measured batch effect removal, reflecting the degree of integration across technologies into a shared embedding space. For each category we computed aggregate scores as the mean of their corresponding metrics.

Lloki achieved the best balance between preserving biological signals and removing batch effects (**Fig**. 2b). It preserved biological variation between cell types, attaining the highest biological conservation score (0.67) and batch correction score (0.85). In contrast, Lloki-FP alone, though yielding a moderate biological conservation score (0.44), performed poorly in batch correction (0.31), highlighting the importance of Lloki-CAE in aligning Lloki-FP embeddings across batches. Among baseline methods, scVI and STAligner achieved the next highest batch correction scores (0.70 and 0.69, respectively), but their biological conservation scores were substantially lower (0.49 and 0.31). While scVI was able to separate major cell types such as excitatory neurons, inhibitory neurons, and oligodendrocytes, it failed to distinguish the remaining cell types. STAligner showed similar limitations, with UMAPs revealing a collapse of the embeddings into a unified structure with minimal cell type separation. This suboptimal integration is likely due to limited gene panel overlap – only 19 shared genes – restricting the ability to fully capture the spectrum of biological variation. PCA embeddings yielded low biological conservation (0.36) and batch correction (0.40), showing weak cell type specificity in the UMAP visualization. Seurat and Harmony improved batch correction relative to PCA (0.62) but further reduced biological conservation (0.35 and 0.31), indicating a trade-off in improved batch alignment at the expense of cell type resolution.

To dissect the contributions of Lloki-CAE’s three-part loss function and to evaluate the importance of the Lloki-FP feature transformation, we conducted two ablation experiments (**Fig**. S1). In the first experiment, we removed each component of the loss function. Removing the biological conservation loss had the most significant impact, reducing the biological conservation score by 0.37, underscoring its critical role in maintaining cell type separation in the embedding space. In contrast, removing the triplet loss compromised batch correction, highlighting the delicate balance required between these objectives. In the second experiment, we compared Lloki-CAE’s performance when using scGPT embeddings versus the transformed Lloki-FP embeddings as input. While using scGPT embeddings alone improved batch correction by 0.08, it biological conservation score dropped by 0.18. Together, these results suggest that while Lloki-FP is highly effective at maintaining biological variation and cell type separation, Lloki-CAE is crucial for integrating slices across technologies and successfully removing batch effects.

### Lloki achieves robust cross-technology spatial alignment

We evaluated Lloki’s ability to align multiple ST slices within a common spatial coordinate framework while ensuring key tissue features are properly aligned. To assess its robustness, we tested two key aspects: (1) its ability to align slices across different technologies, and (2) its performance when slices have minimal overlap in their gene panels. For slice alignment, Lloki embeddings were used to generate landmark pairs between ST slices by identifying mutual nearest neighbors within the Lloki embedding space. These landmark pairs were then aligned using the Kabsch algorithm [29], which finds the optimal rotation and translation to minimize root-mean-square deviation (RMSD) between paired landmarks.

First, we evaluated Lloki’s ability to align slices from different technologies by selecting one slice each from the MERSCOPE, MERFISH, and STARmap datasets (**Fig**. 3a (left)) and performing pairwise alignments using overlapping genes. As baselines, we compared against: (1) the Kabsch algorithm using mutual nearest neighbors from PCA embeddings of shared genes, (2) PASTE [30], a state-of-the-art spatial alignment method that uses fused Gromov–Wasserstein optimal transport to compute alignment based on both transcriptional and spatial similarity, (3) STAligner [7], which provides a slice alignment module in its integration framework, and (4) GPSA [31], which uses deep Gaussian processes to map spatial coordinates between slices into a common coordinate system based on gene expression.

**Figure 3:**
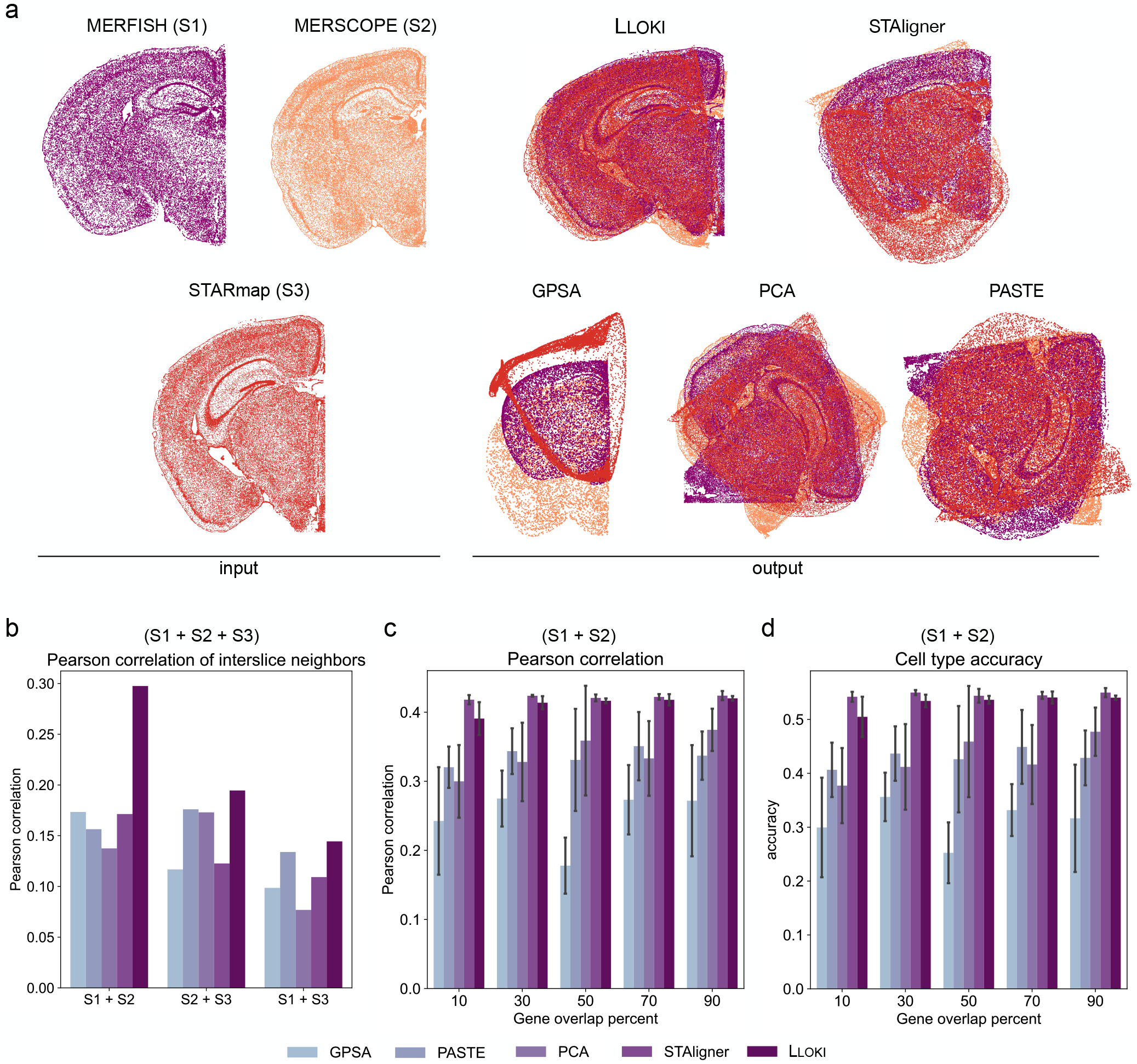
Evaluation of Lloki’s performance in spatial slice alignment. **a.***In situ* visualizations of three slices from the MERFISH, MERSCOPE, and STARmap datasets, along with alignment results using Lloki and four baseline methods. **b**. Alignment accuracy as measured by Pearson correlation of interslice neighbors between each pair of slices from **(a). c-d**. Comparison of Lloki and four baseline methods on alignment accuracy of the MERFISH and MERSCOPE slices as gene overlap decreases, showing results for **(c)** Pearson correlation, and **(d)** cell type accuracy.

All four baselines failed to correctly align the slices, however, Lloki successfully aligned them such that corresponding regions overlapped (**Fig**. 3a (right)). PCA, PASTE, and STAligner all rotate at least one of the slices incorrectly. In addition to scaling, rotating, and translating the slices, GPSA additionally allows non-linear scaling, which can produce artifacts like those seen in their alignment when batch effect is pronounced.

To quantify alignment accuracy, we analyzed each pair of aligned slices by defining interslice cell pairs, where each cell in one slice was paired with its nearest spatial neighbor from the other slice in the aligned coordinate space. For each pair, we computed Pearson correlation using their shared genes. Lloki outperformed all baselines across all three slice pairs (**Fig**. 3b). In particular, for the MERFISH– MERSCOPE pair, Lloki achieved a Pearson correlation of 0.30, compared to 0.17 for the next-best methods, GPSA and STAligner.

To test Lloki’s performance under low gene panel overlap, we used the MERFISH and MERSCOPE slices and created ten simulated datasets by progressively downsampling their shared genes in 10% increments, repeating this process five times per increment to create replicates. Because these two datasets share unified cell type annotations, we also measured how often interslice pairs share the same annotated cell type, in addition to computing Pearson correlation. Before alignment, we applied a random rotation and translation to one slice to ensure that alignment methods were actively aligning the data, rather than relying on preexisting spatial similarity.

Lloki consistently outperformed GPSA, PASTE, and PCA, and performed comparably to STAligner (**Figs**. 3c and 3d). Notably, Lloki and STAligner maintained stable performance with few overlapping genes in this pairwise cross-technology alignment, while the other methods exhibited decreased performance.

Overall, these results demonstrate that Lloki embeddings are highly robust to differences in gene panels, enabling accurate alignment even with minimal overlap between slices. Additionally, Lloki effectively mitigates batch effects across technologies, facilitating accurate cross-technology slice alignment for datasets from diverse ST platforms.

### Lloki enhances cross-technology spatial gene program detection

To assess Lloki’s utility for downstream analysis, we integrated data from four ST platforms – MER-FISH, MERSCOPE, CosMx, and Xenium – and evaluated whether Lloki embeddings improve cross-technology analyses using SpiceMix [32]. SpiceMix models gene expression as a combination of latent factors (“metagenes”) that capture transcriptional programs with spatial organization. Joint analysis enables a more comprehensive understanding of tissue biology by revealing conserved spatial patterns that may be missed when datasets are analyzed in isolation. However, SpiceMix requires a common feature space, which limits its application to datasets with few overlapping genes. We address this limitation using Lloki embeddings as an alternative input to SpiceMix, providing a unified feature space across datasets.

We computed metagenes using SpiceMix under three conditions: (1) Lloki-based integration, using 128-dimensional Lloki embeddings as input to SpiceMix, (2) a shared-gene approach, applying SpiceMix to the 22 genes common across datasets, and (3) independent SpiceMix runs on each dataset, with a post hoc optimal metagene mapping.

To evaluate the consistency and biological relevance of metagenes across conditions, we matched metagenes to the major cell types annotated in each dataset (**Fig**. 4a). For the Lloki-based integration, we found consistent associations between metagenes and major cell types across datasets. In particular, metagene 1 was enriched in astrocytes, metagene 6 in microglia, metagenes 5, 7, and 14 in vascular cells, and metagenes 11 and 13 in oligodendrocytes, with additional metagenes capturing excitatory and inhibitory neurons. In contrast, both the shared-gene and independent approaches captured only partial associations, failing to consistently identify microglia- and oligodendrocyte-specific metagenes.

**Figure 4:**
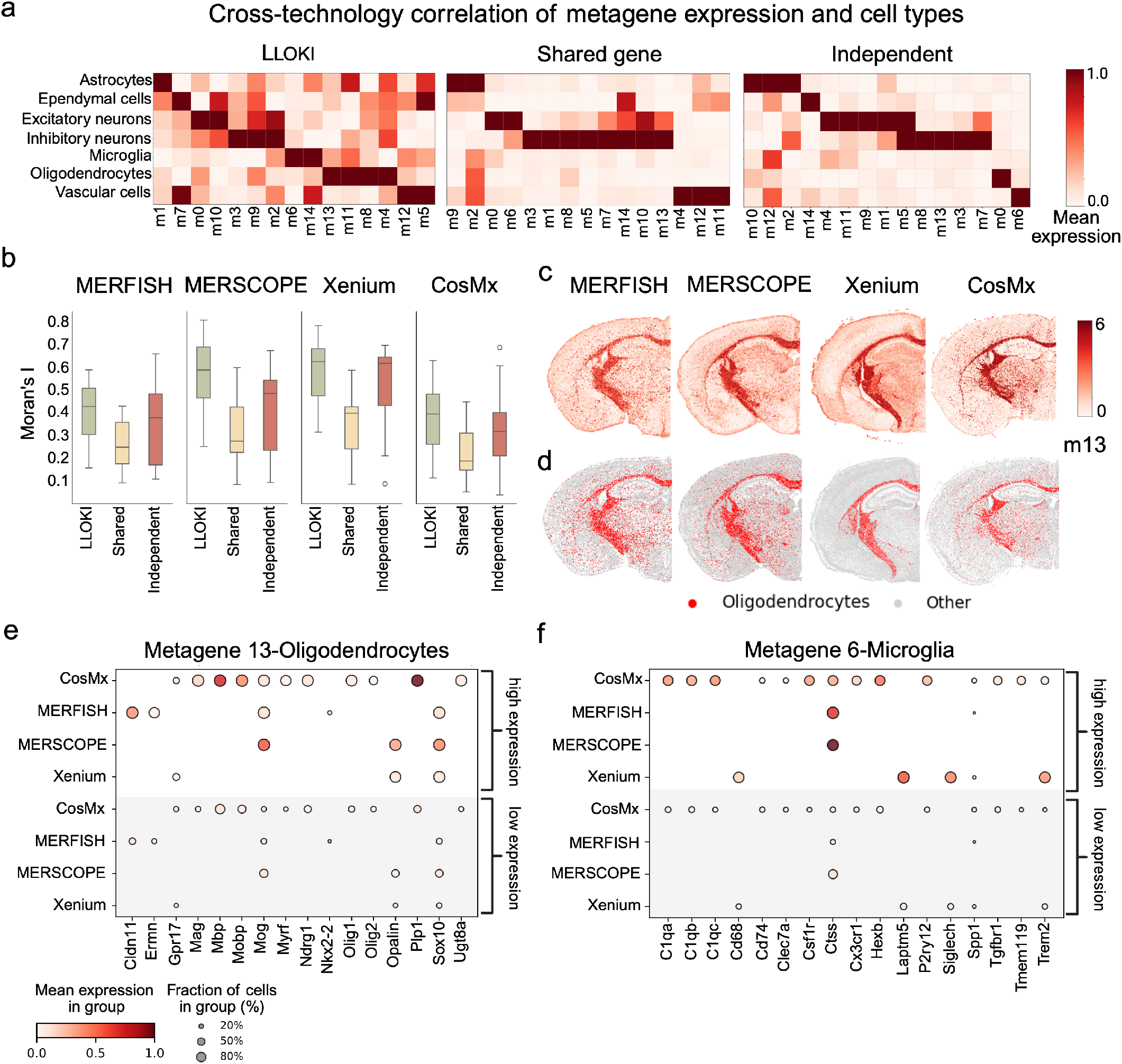
SpiceMix with Lloki embeddings enhances cross-technology metagene discovery. **a.**Cell type enrichment of metagenes across MERFISH, MERSCOPE, CosMx, and Xenium datasets, comparing three approaches: Lloki-based integration, the shared-gene approach (22 common genes), and independent analyses. **b**. Quantitative assessment of metagene quality using Moran’s I spatial autocorrelation scores, shown as box plots. **c**. Spatial distribution of metagene 13 across all four technologies, showing consistent patterns using Lloki embeddings with SpiceMix. **d**. Spatial distribution of oligodendrocytes across MERFISH, MER-SCOPE, Xenium, and CosMx, demonstrating the correspondence between metagene 13 and oligodendrocytes. **e**. Expression levels of canonical oligodendrocyte marker genes across MERFISH, MERSCOPE, Xenium, and CosMx in cells highly expressing metagene 13 versus cells lowly expressing metagene 13. **f**. Expression levels of canonical microglia marker genes across MERFISH, MERSCOPE, Xenium, and CosMx in cells highly expressing metagene 6 versus cells lowly expressing metagene 6.

We further evaluated metagene quality for each run using Moran’s I to quantify spatial autocorrelation (**Fig**. 4b). In brain tissue, functionally similar cells tend to cluster spatially, so a metagene exhibiting spatial autocorrelation indicates that it recapitulates known cell-type patterns. Our results show that the Lloki-based approach consistently produced metagenes with strong spatial autocorrelation in all datasets, while the shared-gene approach exhibited the lowest autocorrelation, and the independent analyses showed high variability.

To assess biological relevance, we examined metagenes with consistent spatial and cell-type associations in all datasets. We found that metagene 13 showed consistent spatial expression patterns in all four datasets (**Fig**. 4c) and was enriched in oligodendrocytes (**Fig**. 4d), indicating it captures a cell type-specific gene program.

To further investigate the molecular underpinnings of metagene 13, we stratified cells by metagene expression and cross-referenced their original gene panels to compare expression of oligodendrocyte marker genes between groups (**Fig**. 4e). Nearly all oligodendrocyte markers exhibited strong correlation with metagene 13. Many of these markers were only measured in one dataset, including *Mbp, Mobp*, and *Plp1* in CosMx, and *Cldn11* and *Ermm* in MERFISH. Other markers, such as *Mog, Opalin*, and *Sox10*, were measured in multiple datasets, Collectively, these findings highlight the value of leveraging full gene panels via Lloki – rather than restricting analysis to a limited set of shared genes – and validate the association of metagene 13 with oligodendrocyte identity.

We used a similar approach to associate metagene 6 with microglia (**Fig**. 4f). In both MERFISH and MERSCOPE, cells with high metagene 6 expression consistently exhibited elevated expression of the microglia marker *Ctss*. Higher expression of metagene 6 was associated with increased expression of *Laptm5, Siglech*, and *Trem2* in Xenium, and *C1qa, C1qb, C1qc*, and *Hexb* in CosMx. These markers – many of which are uniquely expressed across the different technologies – enabled robust identification of a conserved program in microglia only when using full gene panels with Lloki. As with metagene 13, microglia-associated metagenes were not consistently detected using baseline methods.

Together, these results show that Lloki’s integrated embeddings enable SpiceMix to delineate biologically meaningful spatial gene programs, overcoming the limitations imposed by dataset-specific gene panels and enhancing cross-technology analyses. For further validation, *in situ* metagene plots for SpiceMix runs using Lloki embeddings, shared-genes, and independent gene panels are provided in **Figs**. S2, S3, and S4, respectively.

### Lloki identifies shared gene program for tumor infiltration in ovarian cancer

We next applied Lloki to integrate five ST datasets of human ovarian cancer [33], which differ in both gene panel composition and technology (**Fig**. 5a). These datasets include three from CosMx (SMI), one from Xenium (ISS), and one from MERFISH. The CosMx datasets include: (1) 960 genes measured across 100 small tissue samples (0.9 mm × 0.6 mm), (2) 1,000 genes measured across 4 large whole-tissue samples (up to 2 cm), and (3) 6,175 genes measured across 62 small tissue samples (0.5 mm × 0.5 mm). The Xenium dataset contains 240 genes from 32 small tissue samples (1.5 mm × 1.5 mm), while the MERFISH dataset includes 140 genes from 4 large whole-tissue samples (up to 2 cm). Integration via Lloki achived robust mixing of shared cell types across these technologies (**Fig**. 5b), yielding a biological conservation score of 0.47 and a batch correction score of 0.23 when evaluated on non-malignant cells. Malignant cells remained largely distinct between datasets, reflecting tumor-specific transcriptional differences from diverse origins.

**Figure 5:**
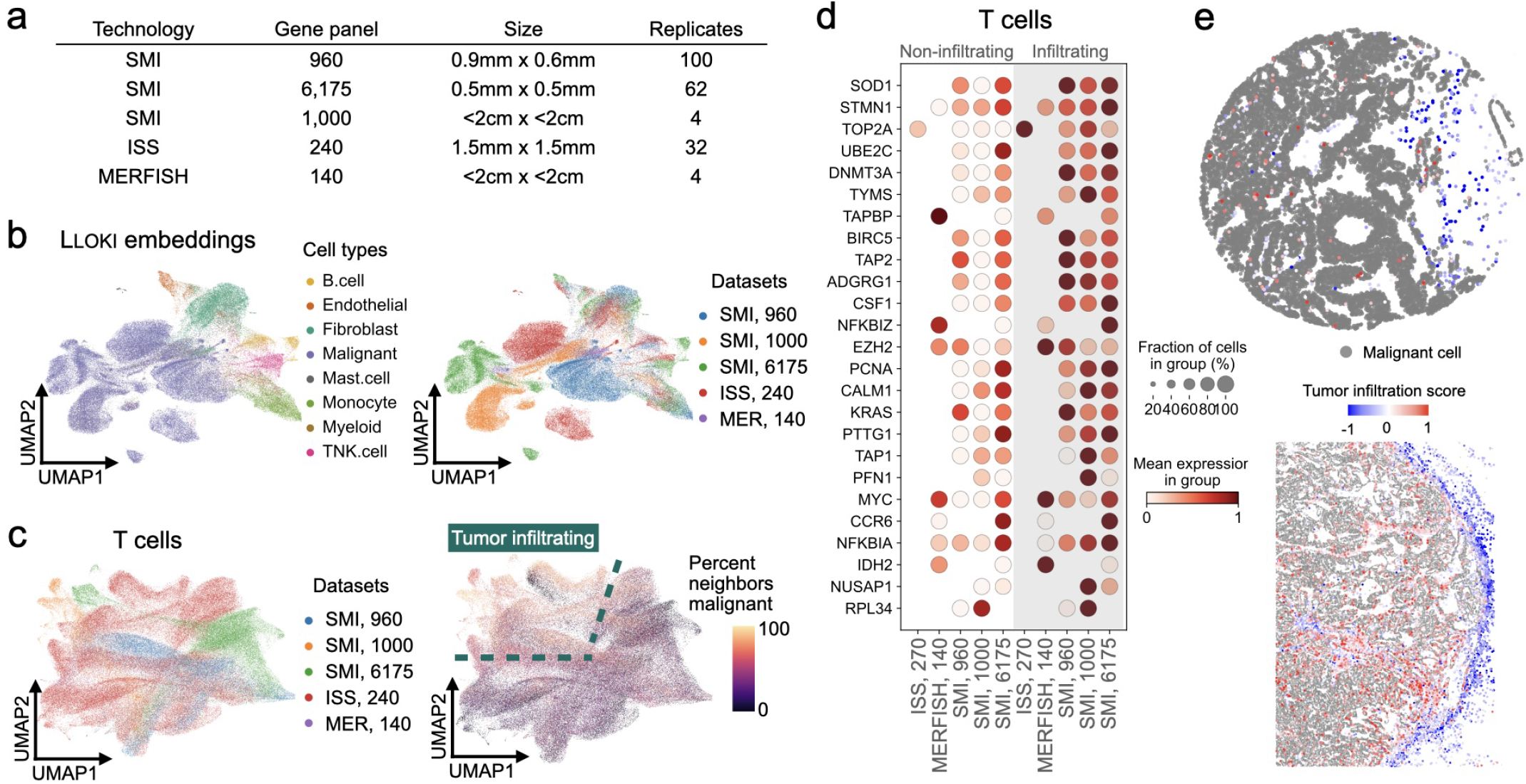
Lloki identifies cross-technology gene program indicative of tumor infiltration in T cells in ovarian cancer. **a.**Summary of the five ovarian cancer ST datasets used in this analysis. **b**. UMAP visualization of Lloki embeddings shows successful integration of non-malignant cells across datasets, while transcriptionally distinct tumor cells remain separate. **c**. UMAP visualization of Lloki embeddings of T cells, colored by dataset (left) or by the proportion of 100-nearest spatial neighbors that are malignant. Dashed outlines indicate regions identified as tumor-infiltrating. **d**. Expression levels of upregulated genes from the cross-technology tumor infiltration gene program in infiltrating vs. non-infiltrating T cells. **e**. Representative *in situ* visualization showing T cells colored by the tumor infiltration score and malignant cells in gray.

Using Lloki embeddings, we further investigated T cell heterogeneity with a focus on tumor infiltration. We combined T cells from all datasets and clustered them based solely on their Lloki embeddings. This unsupervised approach partitioned T cells into two groups that corresponded well with the percentage of malignant cells among their 100 nearest spatial neighbors – a proxy for local tumor infiltration (**Fig**. 5c).

To characterize the genes responsible for this tumor infiltration program, we performed integrated differential expression analysis in two stages: (1) comparing T cells against all other cell types to obtain T cell-specific genes for each dataset’s gene panel, and (2) identifying differentially expressed genes between the two T cell groups detected via Lloki embeddings. Genes consistently up- or downregulated across datasets were retained, yielding a shared gene panel that reliably distinguishes tumor-infiltrating T cells (**Fig**. 5d). In particular, this gene panel includes markers associated with cell proliferation (e.g., *STMN1, TOP2A, UBE2C, BIRC5, NUSAP1*) and antigen processing (e.g., *TAP1, TAP2, TAPBP*), suggesting that these processes are correlated with tumor infiltration.

To assess the biological relevance of this gene panel, we devised a tumor infiltration score, weighting upregulated genes positively and downregulated genes negatively. When spatially projected onto individual T cells, this score revealed a clear pattern: T cells within tumor regions exhibited consistently higher scores than those outside the tumor (**Fig**. 5e). We further quantified the correlation between this tumor infiltration score and the percentage of malignant cells among the 100 nearest spatial neighbors. Across 170,000 T cells from all datasets, we observed a significant correlation (Pearson *R* = 0.326, *P <* 1 × 10^−200^; Spearman *ρ* = 0.366, *P <* 1 × 10^−200^), confirming the biological relevance of our scoring approach.

These findings further demonstrate that Lloki not only enables effective integration of heterogeneous spatial transcriptomics data but also facilitates the discovery of biologically meaningful gene programs, such as the one defining tumor-infiltrating T cells in ovarian cancer.

## Discussion

In this work, we developed Lloki, a novel framework for integrating ST data across diverse platforms without requiring shared gene panels. Lloki combines feature alignment and batch integration through two complementary components: Lloki-FP, which aligns gene panels into a shared feature space by addressing differences in data sparsity, and Lloki-CAE, which mitigates batch effects across ST technologies while preserving biological specificity. By leveraging both components, Lloki enables robust cross-technology integration while retaining meaningful biological variation.

Our results demonstrate that Lloki outperforms baseline methods across key integration metrics, maintaining biological variation and effectively correcting batch effects – even when gene panel over-lap is minimal. Lloki’s integrated embeddings support challenging downstream tasks such as spatial gene program identification and slice alignment, underscoring its versatility in addressing the inherent complexities of ST data integration and paving the way for more comprehensive analysis of tissue architecture across diverse technologies.

Several avenues exist for extending Lloki’s capabilities. Refinements to Lloki-FP – such as improved mitigation of technology-specific biases via optimal transport – could enhance the alignment between ST and scRNA-seq data. Similarly, the conditional autoencoder design of Lloki-CAE offers promising generalization capabilities: by updating only the conditional weights while retaining shared parameters, Lloki can be readily adapted to new ST platforms without full retraining. As the field of scFMs evolves, our approach can be extended to incorporate emerging models, further enhancing integration performance.

Beyond methodological advancements, Lloki has several practical applications. One notable use case is leveraging samples from one ST technology as controls for disease samples collected using a different technology, reducing data requirements and conserving resources. Moreover, Lloki could be applied to the integration of atlas-scale datasets, enabling large-scale, cross-technology analyses to identify rare cellular niches without necessitating prohibitively large sample sizes. Overall, Lloki provides a powerful tool for harmonizing ST datasets across different platforms and gene panels, facilitating a deeper understanding of spatial cellular organization in diverse biological contexts.

By addressing key challenges in ST data integration, Lloki lays the groundwork for scalable, cross-technology spatial transcriptomics analysis, empowering researchers to uncover biologically meaningful patterns across diverse tissue samples and experimental conditions.

## Methods

Lloki is a framework for integrating spatial transcriptomics (ST) data from diverse platforms by addressing two key alignment challenges (**Fig**. 1): (1) feature alignment across varying gene panels, handled by Lloki-FP, which imputes gene expression and embeds cells using a single-cell foundation model (scFM); and (2) batch alignment across different ST technologies, handled by Lloki-CAE via a conditional autoencoder with a novel three-part loss function. This approach allows for unified analysis of heterogeneous ST datasets while preserving essential biological signals.

### Feature alignment and denoising with Lloki-FP

scFMs offer a promising solution for aligning features across ST datasets without requiring shared gene panels [11–13]. However, ST data often appears out-of-distribution for scFMs pretrained on scRNA-seq due to differences in sparsity and gene detection sensitivity. Additionally, scFMs process only genes with non-zero expression, making higher sparsity a major challenge, as it reduces the effective feature dimensionality and introduces variability in embedding quality across ST platforms.

To address this, Lloki-FP employs a sparsity-matching and denoising approach, combining Wasserstein optimal transport [34] with graph-based feature propagation [35]. Rather than performing conventional gene imputation, Lloki-FP transforms the sparsity profile of ST data to better match the scRNA-seq distribution, optimizing compatibility with scFMs and improving embedding quality. Unlike prior feature propagation methods for single-cell data [36], Lloki-FP integrates spatial information and computes a sparsity prior via optimal transport, shifting the focus from mere imputation to distribution alignment.

At a high level, Lloki-FP first determines a target sparsity for each ST cell using optimal transport. It then constructs a cellular graph incorporating both gene similarity and spatial proximity, which is used to iteratively impute missing gene expression until each cell’s target sparsity is reached.

#### Graph Construction

Lloki-FP begins by constructing an undirected graph **G** = (**V, E**), where nodes *v*_*i*_ ∈ **V** represent cells, and edges *e*_*ij*_ ∈ **E** encode similarity between cells. Since feature propagation assumes high graph homophily – where connected cells share similar gene expression – it is critical that the graph reflects true biological similarity.

To achieve this, Lloki-FP integrates gene expression and spatial proximity into graph construction. Each cell’s *k*-nearest neighbors (*k*-NNs) are identified in the gene expression space to establish graph connectivity, while edge weights (*w* : *E* → **ℛ**^+^) are set using a Radial Basis Function kernel based on spatial distance:

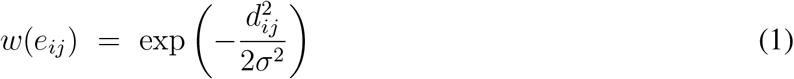

where *d*_*ij*_ is the spatial distance between cells *i* and *j* and *σ* is a bandwidth parameter computed as the mean distance from a random sample of 100 cell pairs. This weighting reinforces graph homophily by leveraging the biological insight that physically adjacent cells tend to share microenvironments and gene expression patterns. The graph is then symmetrically normalized to ensure balanced feature propagation across cells. This graph-guided feature propagation aligns ST sparsity distributions with scRNA-seq while ensuring imputation is informed by both transcriptional and spatial information.

#### Feature propagation for data imputation with optimal transport

To impute sparse gene expression values, Lloki-FP employs feature propagation [35], enhanced with optimal transport [34] to align ST sparsity with scRNA-seq references. Given that ST data is typically more sparse than scRNA-seq data, our approach selectively imputes missing values while preserving a degree of inherent sparsity – since completely eliminating sparsity would reduce embedding diversity and obscure biologically meaningful variation.

This process starts with the constructed similarity graph **G** with spatially informed, normalized adjacency matrix **A** and the original gene expression matrix **X**. Gene expression is iteratively updated as follows:

1. *Feature propagation with controlled sparsity*. Imputed gene expression values are updated iteratively as:

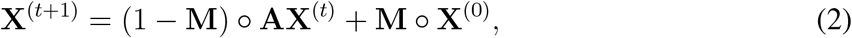

where **X**^(0)^ is the original gene expression matrix, **M** is a binary mask indicating observed (nonzero) entries in **X**^(0)^, and ◦ denotes the element-wise Hadamard product. This update minimizes the graph Dirichlet energy as shown in [35], ensuring that imputation is both spatially coherent and biologically relevant.
2. *Sparsity matching with optimal transport*. To harmonize sparsity profiles between ST and scRNA-seq data, we compute per-cell sparsity as 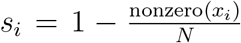, where nonzero(*x*_*i*_) is the number of nonzero gene expression values in cell *i* and *N* is the total number of genes. We then compute the empirical cumulative distribution function (CDF) for the sparsity values. For a sorted set {*s*_(1)_, *s*_(2)_, …, *s*_(*n*)_}, the CDF is defined as:

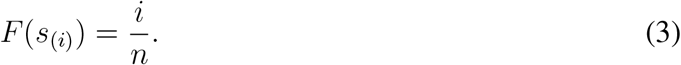 Denoting the CDFs for the ST data and the scRNA-seq reference as *F*_ST_(*s*) and *F*_scRNA_(*s*) respectively, we align the distributions by mapping each ST sparsity value *s* to a target value *s*^*^ via:

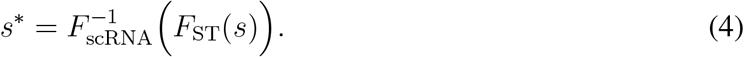 This quantile alignment minimizes the Wasserstein distance between the distributions, yielding target sparsity levels that mirror the scRNA-seq profile. Subsequently, the same feature propagation update is applied iteratively until each cell’s imputed expression conforms to its target sparsity. To prevent over-imputation – where the imputed data becomes denser than intended – we monitor the number of nonzero entries per cell and reintroduce zeros as needed to recapitulate the natural dropout observed in ST technologies.

This iterative process – alternating between controlled feature propagation and sparsity adjustment via 1D optimal transport mapping – ensures that Lloki-FP embeddings preserve critical biological signals while aligning with scRNA-seq sparsity, thereby enhancing compatibility with scFMs.

#### Denoising with feature diffusion

Following optimal transport-enhanced feature propagation, we refine gene expression data using feature diffusion. We construct a new cell similarity graph based on the imputed gene expression matrix **X**_imputed_, with **A**_imputed_ as the corresponding adjacency matrix. During this phase, the original gene expression matrix is diffused across the new graph without modifying non-zero values, using:

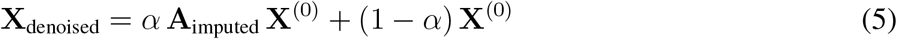

This diffusion step propagates gene expression features across the spatially weighted *k*-NN graph, updating missing or noisy values while preserving the original gene expression measurements. The hyper-parameter *α* ∈ [0, 1] controls the extent to which information from neighboring cells (via **A**_imputed_ **X**^(0)^) influences the denoised output. A value of *α* close to 1 emphasizes the diffused signal from neighboring cells, while a value closer to 0 favors retaining the original gene expression values. This formulation leverages the assumption that cells that are both spatially and transcriptionally similar share comparable true gene expression patterns.

#### Feature alignment using a single-cell foundation model

Once ST sparsity is aligned to match the scRNA-seq reference and the data is denoised, we proceed with feature alignment via a scFM pretrained on scRNA-seq data. This step ensures that all datasets regardless of their original gene panels share a common feature space, allowing the full gene panel to inform cell embeddings.

scGPT [13] demonstrated strong performance across cell type clustering metrics (**Supplemental Information**). It is not only computationally efficient, but also particularly adept at handling smaller gene panels due to its autoregressive pretraining strategy. The final output of Lloki-FP is a 512-dimensional embedding space, consistent across all input datasets, enabling robust feature alignment across diverse ST technologies.

We compare the performance of Lloki-FP to scGPT alone (without feature propagation) across five ST technologies (**Supplemental Information**). Lloki-FP consistently outperforms scGPT, with the most significant improvement observed on the STARmap dataset (ARI of 0.423 compared to 0.197). We also conducted an ablation study evaluating the impact of the optimal transport sparsity alignment step in Lloki-FP (**Supplemental Information**). This analysis revealed that incorporating optimal transport alignment significantly enhanced performance for the MERSCOPE and STARmap datasets, confirming the importance of this step in our workflow.

### Batch integration with Lloki-CAE

Cell embeddings from Lloki-FP effectively capture biological variation within each ST dataset, but still exhibit large batch effects between technologies. To address this, we develop a conditional autoencoder (CAE) for batch integration, conditioned on both ST technology and specific gene panel, which allows the model to differentiate between datasets by technology or gene panel. Integration is achieved through a novel three-part loss function that aligns embeddings across technologies while preserving the structure of cell-type clusters within each dataset.

#### Conditional autoencoder architecture

Our CAE architecture follows a standard design with some key new developments. Both the encoder and decoder comprise three layers, reducing the dimensionality of the 512-dimensional input embeddings from Lloki-FP down to 128. Many downstream tasks require positive embeddings, so we apply a softplus activation function in the final layer of the encoder to ensure all embeddings are positive.

The conditional component of the autoencoder is implemented as a learnable embedding that maps a batch token to a 10-dimensional embedding, appended to the input of both the encoder and decoder.

#### Three-part loss function

Conventional methods for integrating ST datasets – such as STAligner [7] – typically balance cell-type heterogeneity and batch integration using a combination of reconstruction loss and triplet loss. Although triplet loss effectively promotes batch integration, we found that reconstruction loss alone does not sufficiently preserve critical biological information. We therefore introduce a third loss function, the biological conservation loss, designed to maintain the local structure of the original Lloki-FP embeddings, which inherently capture robust biological signal across cell types. By jointly optimizing reconstruction loss, triplet loss, and our new biological conservation loss, our approach aligns embeddings across batches while preserving essential biological information.

1. **Reconstruction loss**. The reconstruction loss seeks to recreate the original Lloki-FP embeddings for each cell *X*. Given the decoder output *Y*, we minimize the mean squared error (MSE) between *X* and *Y*, ensuring that the CAE preserves original information through the encoding-decoding process:

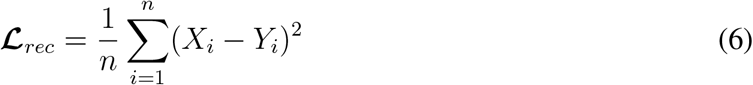
2. **Triplet loss**. Triplet loss encourages embeddings of biologically similar cells from different technologies to be closer, while pushing apart embeddings of dissimilar cells within the same technology. Anchor-positive pairs consist of a cell from one technology (anchor) and its mutual nearest neighbor from another. Negative pairs are randomly selected cells from the same slice as the anchor, which are likely to be dissimilar. When reliable cell type annotations are available, we optionally refine this selection by: (1) restricting positive pairs to cells sharing the same cell type across batches, and (2) selecting negative pairs using hard negative sampling, where the negative pair is chosen as the closest cell from the same batch that belongs to a different cell type than the anchor. The triplet loss is defined as:

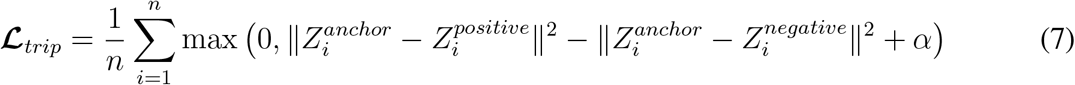

where *Z*_*i*_ is the CAE embeddings, and *α* is the margin separating positive and negative pairs.
3. **Biological conservation loss**. Our new biological conservation loss preserves cell-to-cell similarity by ensuring that the distances between each cell and its *k*-nearest neighbors in the original Lloki-FP embedding space are maintained in the Lloki-CAE embedding space. The biological conservation loss is defined as:

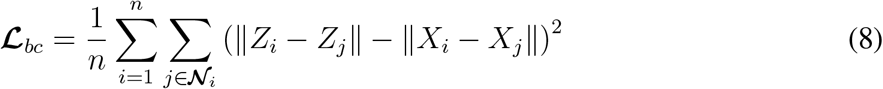

where **𝒩**_*i*_ is the set of nearest neighbors of cell *i* in the original embedding space.

To balance these components, we introduce weighting parameters *λ*_*rec*_, *λ*_*trip*_, and *λ*_*bc*_, which are tuned to determine the optimal balance among the three competing objectives. The total loss is defined as:

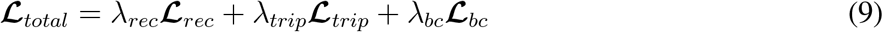

These weights are optimized to ensure that each aspect of the loss appropriately influences training, preserving cell-type clustering while effectively reducing batch effects across technologies.

Further details about hyperparameter choices for Lloki-CAE are in **Supplemental Information**.

### Evaluation metrics for batch integration

To quantitatively evaluate Lloki and compare it to baselines for batch integration we used eight metrics from scib-metrics [28]. Five of these metrics assessed the biological variation preserved by the embeddings, based on cell type separation using cell type annotations provided with each dataset: isolated labels silhouette score, *k*-means clustering NMI and ARI (KMeans NMI and KMeans ARI), silhouette width (Silhouette label), and cell-type local inverse Simpson index (cLISI). The remaining three metrics evaluated batch effect removal, measuring how well technologies were integrated into a shared embedding space: silhouette batch score, batch connectivity in a cell-type specific subgraph (graph connectivity), and principal component regression comparison (PCR comparison). Aggregate scores were computed by averaging biological conservation and batch correction metrics.

### Scalability of Lloki

Lloki demonstrates robust scalability for processing large ST datasets. The computational framework consists of two main components with complementary performance characteristics. Lloki-FP processes each ST slice independently, resulting in computational requirements that scale linearly with the number of slices. For a typical slice containing around 50,000 cells, running Lloki-FP takes approximately 5 minutes and uses 20 GB of memory. Importantly, this component can be fully parallelized across compute nodes to process multiple slices simultaneously, offering substantial acceleration for large-scale studies. Lloki-CAE employs an efficient batching approach during training and maintains a lightweight memory footprint of less than 2 GB regardless of dataset size. For the five-slice integration task described in our results, model training was completed in under one hour on a single GPU. Post-training embedding generation is computationally efficient, requiring only a single forward pass operation through the trained encoder. This process takes less than one second to generate embeddings for cells from all five slices (250,000 cells), enabling rapid integration of new datasets without retraining. Notably, only one slice of each technology is needed to train the model; processing all slices from large datasets after training takes only 5 minutes each and is fully parallelizable.

### Slice alignment

For slice alignment with Lloki, we implemented the Kabsch algorithm [29] for image alignment and used our Lloki embeddings to determine landmark pairs between two slices. The algorithm then seeks to minimize the root-mean-squared distance (RMSD) for all landmark pairs. For baselines, we ran PASTE, STAligner, GPSA, and a version of the Kabsch algorithm that uses PCA embeddings of shared gene expression instead of Lloki embeddings. We ran PASTE with default parameters and five different values for alpha (0.01, 0.05, 0.001, 0.005, 0.0001), using the best-performing parameter set for our comparison. We note that we run PASTE instead of PASTE2 [37], because the advantage of PASTE2 is in partial alignment, and all of our analysis was on full-slice alignments. We ran STAligner and GPSA using their default parameters.

### SpiceMix analysis

For the SpiceMix analysis on the four mouse brain coronal slices from MERFISH, MERSCOPE, Xenium, and CosMx, we ran SpiceMix with the following parameter settings. We set the number of metagenes *K* = 15 and the spatial affinity regularization to 10^−4^. We performed pretraining without optimizing spatial affinity parameters for 10 iterations, followed by 200 iterations with full optimization. For baseline comparisons where SpiceMix was run independently on each dataset using full gene panels, we constructed a cross-dataset metagene mapping to enable direct comparison of metagene identities across technologies. To do this, we first computed metagene signature matrices for each dataset, defined as the Pearson correlation between each metagene and the unified cell type annotations (excluding “Other/Unannotated” cells). These signatures capture the degree to which each metagene is associated with distinct cell types. For each pair of datasets, we computed metagene similarity based on their signature vectors and applied the Hungarian algorithm [38] to derive a one-to-one metagene mapping that maximizes correspondence. Using MERSCOPE as the reference, we identified the best-matched metagene in each of the other datasets (MERFISH, Xenium, and CosMx) for each reference metagene and retained only those with consistent matches in at least two additional datasets. This mapping was then used to reorder the metagene dimensions in the independently trained datasets’ SpiceMix embedding matrices, so that mapped metagenes align across datasets.

To assess the correspondence of metagenes identified by SpiceMix with cell type annotations, we computed an enrichment matrix where each row represents a cell type and each column a metagene, using only high-confidence cell type annotations. For each dataset, we excluded “Other/Unannotated” cells, retaining seven cell type labels. For each dataset, we computed the mean metagene expression for each cell type using only non-zero values, then aggregated the results into a unified matrix. Metagenes were then reordered based on their maximum enrichment across cell types to highlight cell type-specific patterns and enable cross-technology comparisons.

We used Moran’s I to evaluate spatial autocorrelation of metagene expression for each dataset. Spatial neighborhood graphs were constructed using Squidpy [39]. For each metagene, Moran’s I was computed to quantify how strongly its expression was spatially clustered.

To further evaluate the biological relevance of individual metagenes, we conducted a marker gene analysis stratified by metagene expression. Specifically, for metagenes of interest (e.g., metagene 13 for oligodendrocytes, metagene 6 for microglia), we divided cells into high and low expression groups based on the 95th percentile of metagene expression within each dataset. We then examined the expression of known cell type–specific marker genes across these groups using the original, unprocessed gene expression matrices. For each gene and group, we computed the mean expression and the fraction of expressing cells (non-zero expression), enabling comparison across technologies with non-overlapping gene panels.

### Ovarian cancer analysis

We integrated the ovarian cancer datasets using Lloki with the following hyperparameters: *λ*_*bc*_ = 500.0, *λ*_*trip*_ = 0.2, *λ*_*recon*_ = 1.0, a learning rate of 0.0005, a chunk size of 16,000, and a batch dimension of Hyperparameter tuning was performed to optimize for both a high biological conservation score and effective batch correction which was measured using scib-metrics.

To identify tumor infiltrating T cells, we first performed Leiden clustering on T cells from all datasets, using Lloki embeddings. We then selected clusters corresponding to regions in the UMAP with a high percentage of malignant neighbors, thereby defining a cross-technology group of tumor-infiltrating T cells.

To derive a gene list associated with tumor infiltration, we used a two-stage process. In the first stage, for each dataset, we performed differential expression analysis comparing T cell populations to identify genes that were enriched in T cells. We applied filters to retain only genes expressed in at least 5% of T cells and exhibiting a minimum fold change of 0.3. This produced 816 T cell-enriched genes in all datasets. In the second stage, we split T cells by infiltration status and repeated differential expression analysis on the subset of genes identified in the first stage. These results were aggregated using a weighted rank-based aggregation method: for each dataset, each gene received a normalized ranking and the final ranking was averaged across datasets.

Finally, using the shared gene list, we scored T cells for their expression of the tumor infiltration program with Scanpy, using positive weights for upregulated genes and negative weights for downregulated genes. This provided a robust metric to quantify T cell infiltration in the integrated ovarian cancer datasets.

## Acknowledgment

This work was supported, in part, by National Institutes of Health Common Fund 4D Nucleome Program grant UM1HG011593 (J.M.); National Institutes of Health Common Fund Cellular Senescence Network Program grant UG3CA268202 (J.M.); and National Institutes of Health grants R01HG007352 (J.M.), R01HG012303 (J.M.), R21DA061481 (J.M.), and U24HG012070 (J.M.). J.M. was additionally supported by the Ray and Stephanie Lane Professorship, a Guggenheim Fellowship from the John Simon Guggenheim Memorial Foundation, a Google Research Award, and a Single-Cell Biology Data Insights award from the Chan Zuckerberg Initiative. S.K. is a Lane Fellow. The funders had no role in study design, data collection and analysis, decision to publish or preparation of the manuscript.

## Code Availability

The source code for Lloki can be accessed at: https://github.com/elliehaber07/LLOKI.

## Author Contributions

Conceptualization, E.H., S.K., J.M.; Methodology, E.H., S.K., J.M.; Software, E.H., A.D., S.K.; Investigation, E.H., S.K., J.M.; Writing, E.H., S.K., J.M.; Funding Acquisition, J.M.

## Competing Interests

The authors declare no competing interests.

## Notes

### Competing Interest Statement

The authors have declared no competing interest.

### Summary of Updates

We have updated the Results sections (for Figures 2-4).

