## Supplemental Information for "Unified integration of spatial transcriptomics across platforms"

#### A Supplementary Note

##### Clustering performance of LLOKI-FP

To assess the quality of embeddings generated by LLOKI-FP, we compared them to embeddings produced by directly passing the gene expression matrix through scGPT, which serves as our baseline. We applied Leiden clustering to the embeddings generated by each method to produce clusters for metric evaluation. To ensure the number of clusters matched the ground truth, we performed an interval search to determine the resolution parameter that yielded the exact number of clusters as the ground truth. **Table** S1 shows the clustering performance using ARI and NMI scores. As shown, LLOKI-FP consistently outperforms the baseline across all datasets and metrics, demonstrating the effectiveness of our approach.

##### Hyperparameter choice for LLOKI-FP and LLOKI-CAE

Here we provide additional details on training LLOKI-FP and LLOKI-CAE, including hyperparameter selection and optimizations for training efficiency and performance.

For LLOKI-FP, we constructed $k$-NN graphs using $k = 40$ neighbors. As detailed in the **Methods** section, we employed the optimal transport stopping criterion to determine the number of feature propagation iterations. For the feature diffusion step, we empirically determined $\alpha = 0.05$ (self-weight = 0.95) as the optimal value for updating gene expression, and used only a single graph iteration.

LLOKI-CAE's loss function includes three distinct terms, each weighted by a corresponding parameter. The best results were obtained using $\lambda_{\text{rec}} = 1$, $\lambda_{\text{nn}} = 500$, and $\lambda_{\text{trip}} = 2$.

During early training, positive-anchor pairs selected for triplet loss may be suboptimal. To address this, we introduced a warm-up parameter $w$ for the triplet weight. For the first $w$ iterations, the triplet weight $\lambda'_{\text{trip}}$ is scaled as follows:

$$\lambda'_{\text{trip}} := \min\left(\lambda_{\text{trip}}, \lambda_{\text{trip}} \times \frac{i - w + 1}{t}\right)$$

where $t$ is the total number of training iterations and $i$ is the current iteration. We used $w = 10$ for the warm-up parameter. Additionally, triplets were computed after each iteration to ensure continual improvement in the selection of possible triplets.

For the neighborhood loss, we use $k = 30$ neighbors, limited to cells from the same technology. This reduces the neighbor search space and reflects the observation that early in the training, meaningful neighbors are unlikely to come from other technologies.

For the triplet loss, mutual nearest neighbors (MNNs) between each pair of technologies were identified using $k = 40$. All MNNs were retained as anchor-positive pairs, and negative samples were drawn randomly from the same technology as the anchor. For the LLOKI run shown in **Fig.** 2, we utilized the optional cell type refinement described in the **Methods** section. Specifically, rather than using all MNNs, positive pairs were formed by randomly selecting cells from any batch that share the same cell type as the anchor. For negative sampling, instead of random selection, we chose the closest cell from the same batch that belonged to a different cell type. This hard negative sampling strategy improved embedding space differentiation by focusing on the most challenging cases for discrimination.

To manage GPU memory and enable frequent backpropagation, we implemented fixed mini-batching. Batches were computed once at the beginning of training and held constant throughout. This simplification reduces computational overhead and ensures that all neighborhood and triplet computations remain within-batch. We found that a batch size of 16,000 cells provided the best results.

#### Creating a high-level cell type annotation

A key challenge when integrating ST datasets is the variation in available annotations. This variability complicates direct comparisons and the visualization of batch integration performance. To address this, we created a unified set of high-level cell type annotation for use in UMAP visualizations, while retaining the original annotations for evaluation using biological conservation metrics.

To generate this unified annotation, we aggregated the original cell type labels from four datasets (MERSCOPE, MERFISH, STARmap, and CosMx). We mapped these labels into approximately ten broader categories that perserve key biological distinctions. These high-level annotations, along with the original cell types they encompass, are detailed in **Table** S2.

The Xenium dataset did not include cell type annotations but provided cluster identities. To annotate Xenium clusters, we used marker gene expression and spatial localization. Where marker genes were unavailable – for example, for astrocytes – we matched the spatial distribution of Xenium clusters to the known spatial patterns of annotated cell types in other datasets. This heuristic spatial mapping allowed us to assign high-level annotations even in the absence of canonical markers.

### B Supplementary Tables

| | scGPT | | LLOKI-FP | | LLOKI-FP – OT | |
|---|---|---|---|---|---|---|
| **Dataset** | **ARI** | **NMI** | **ARI** | **NMI** | **ARI** | **NMI** |
| MERSCOPE | 0.541 | 0.740 | 0.727 | 0.830 | 0.606 | 0.793 |
| MERFISH | 0.605 | 0.795 | 0.892 | 0.893 | 0.893 | 0.903 |
| STARmap | 0.197 | 0.364 | 0.423 | 0.609 | 0.351 | 0.560 |
| CosMx | 0.347 | 0.555 | 0.395 | 0.599 | 0.370 | 0.607 |
| Xenium | 0.451 | 0.620 | 0.522 | 0.709 | 0.542 | 0.718 |

**Table S1:** Ablation study on cell-type clustering metrics (Adjusted Rand Index, ARI, and Normalized Mutual Information, NMI) comparing LLOKI-FP with scGPT and LLOKI-FP without the optimal transport alignment (*–OT*). Since optimal transport is a key component of LLOKI-FP, removing it enables us to evaluate its specific contribution to clustering performance. Results are shown for five spatial transcriptomics datasets: MERSCOPE, MERFISH, STARmap, CosMx, and Xenium. Higher ARI and NMI values indicate better clustering performance.

| High-Level Annotation | Corresponding Entries |
|---|---|
| Astrocytes | Astrocytes_Cortex_Hippocampus,<br>Astrocytes_Thalamus_Hypothalamus,<br>Astrocytes,<br>Astro-NT,<br>Astro-TE,<br>Astroependymal |
| Ependymal cells | Ependymal cells,<br>Tanycytes,<br>Choroid_plexus_epithelial_cells,<br>Ependymal_cells,<br>Tanycyte,<br>Ependymal,<br>CHOR |
| Excitatory neurons | CNU-HYa Glut,<br>CNU-MGE GABA,<br>Excitatory_neurons_Hippocampal_CA1,<br>Excitatory_neurons_Hippocampal_CA2,<br>Excitatory_neurons_Hippocampal_CA3,<br>Excitatory_neurons_Layer1_Piriform,<br>Excitatory_neurons_Layer2_3,<br>Excitatory_neurons_Layer4,<br>Excitatory_neurons_Layer5,<br>Excitatory_neurons_Layer5_6,<br>Excitatory_neurons_Layer6,<br>Excitatory_neurons_Telencephalon,<br>Peptidergic_neurons,<br>Excitatory_neurons_Amygdala,<br>Excitatory_neurons_Di/mesencephalon,<br>Cholinergic_neurons_Habenebula,<br>HY Glut,<br>HY Gnrh1 Glut,<br>HY MM Glut,<br>IT-ET Glut,<br>MB Glut,<br>MH-LH Glut,<br>MY Glut,<br>NP-CT-L6b Glut,<br>OB-CR Glut,<br>P Glut,<br>Telencephalon projecting excitatory neurons,<br>Di- and mesencephalon excitatory neurons,<br>TH Glut |
| Inhibitory neurons | Cck_interneurons,<br>CNU-HYa GABA,<br>CNU-LGE GABA,<br>CNU-MGE GABA,<br>CTX-CGE GABA,<br>CTX-MGE GABA,<br>Di- and mesencephalon inhibitory neurons,<br>D1_medium_spiny_neurons,<br>D2_medium_spiny_neurons,<br>HY GABA,<br>Inhibitory_interneurons,<br>Inhibitory_neurons_Amygdala,<br>Inhibitory_neurons_Habenula_Hypothalamus,<br>Inhibitory_neurons_Habenula_Thalamus,<br>Inhibitory_neurons_Reticular_nucleus,<br>Interneurons,<br>MB GABA,<br>MY GABA,<br>OB-IMN GABA,<br>Olfactory inhibitory neurons,<br>Peptidergic neurons,<br>Serotonergic_neurons,<br>Telencephalon_inhibitory_neurons,<br>Telencephalon inhibitory interneurons,<br>Telencephalon projecting inhibitory neurons |
| Microglia | Immune,<br>Microglia |
| Oligodendrocytes | Commited_oligodendrocytes,<br>Mature_oligodendrocytes,<br>Oligodendrocyte precursor cells,<br>Myelin_forming_oligodendrocytes,<br>Newly_formed_oligodendrocytes,<br>Oligodendrocytes,<br>Oligodendrocytes_precursor_cells,<br>OPC-Oligo |
| Other/Unannotated | Neuroblasts,<br>Unannotated |
| Vascular cells | Choroid plexus epithelial cells,<br>Vascular,<br>Vascular and leptomeningeal cells,<br>Vascular endothelial cells,<br>Vascular_endothelial_cells,<br>Vascular_leptomeningeal_cells,<br>Vascular_smooth_muscle_cells,<br>Vascular smooth muscle cells,<br>Perivascular_macrophages,<br>Pericytes |

**Table S2:** Unified high-hevel cell type annotations derived, mapped from the four spatial transcriptomics technologies with available cell type annotations: MERFISH, MERSCOPE, STARmap, and CosMx.

### C Supplementary Figures

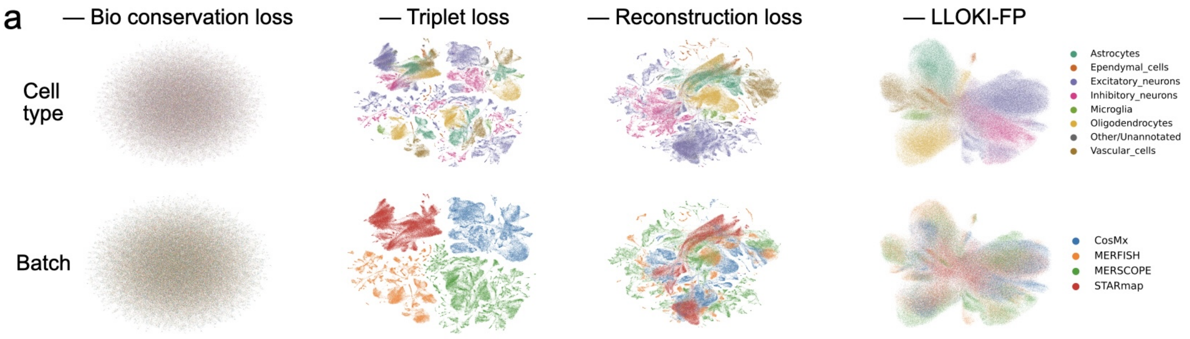


b

| Method | Bio conservation | | | | | Batch correction | | | | Aggregate score | | |
|---|---|---|---|---|---|---|---|---|---|---|---|---|
| | Isolated labels | KMeans NMI | KMeans ARI | Silhouette label | cLISI | Silhouette batch | iLISI | Graph connectivity | PCR comparison | Batch correction | Bio conservation | Total |
| **Normal** | 0.56 | 0.64 | 0.57 | 0.58 | 1.00 | 0.88 | 0.07 | 0.91 | 0.76 | 0.66 | 0.67 | 0.66 |
| **No LLOKIFP** | 0.53 | 0.32 | 0.19 | 0.50 | 0.92 | 0.91 | 0.28 | 0.83 | 0.94 | 0.74 | 0.49 | 0.59 |
| **No AE loss** | 0.53 | 0.32 | 0.22 | 0.49 | 0.98 | 0.83 | 0.06 | 0.87 | 0.87 | 0.66 | 0.51 | 0.57 |
| **No triplet loss** | 0.53 | 0.36 | 0.28 | 0.54 | 1.00 | 0.51 | 0.00 | 0.38 | 0.00 | 0.22 | 0.54 | 0.41 |
| **No BC loss** | 0.50 | 0.00 | 0.00 | 0.50 | 0.52 | 1.00 | 0.75 | 0.20 | 0.00 | 0.49 | 0.30 | 0.38 |

**Figure S1:** Ablation study of LLOKI performance when removing components of the three part loss function of LLOKI-CAE or LLOKI-FP. **(a)** UMAP visualizations for each ablation, with cells colored by cell type (top) and by technology (bottom). Variants include LLOKI-CAE with the biological conservation loss, triplet loss, or reconstruction loss removed, as well as LLOKI-FP trained on raw gene expression passed directly through through scGPT (i.e., without LLOKI-FP. **(b)** Performance comparison of the four ablations using ten metrics, assessing biological variation preservation and batch mixing across technologies.

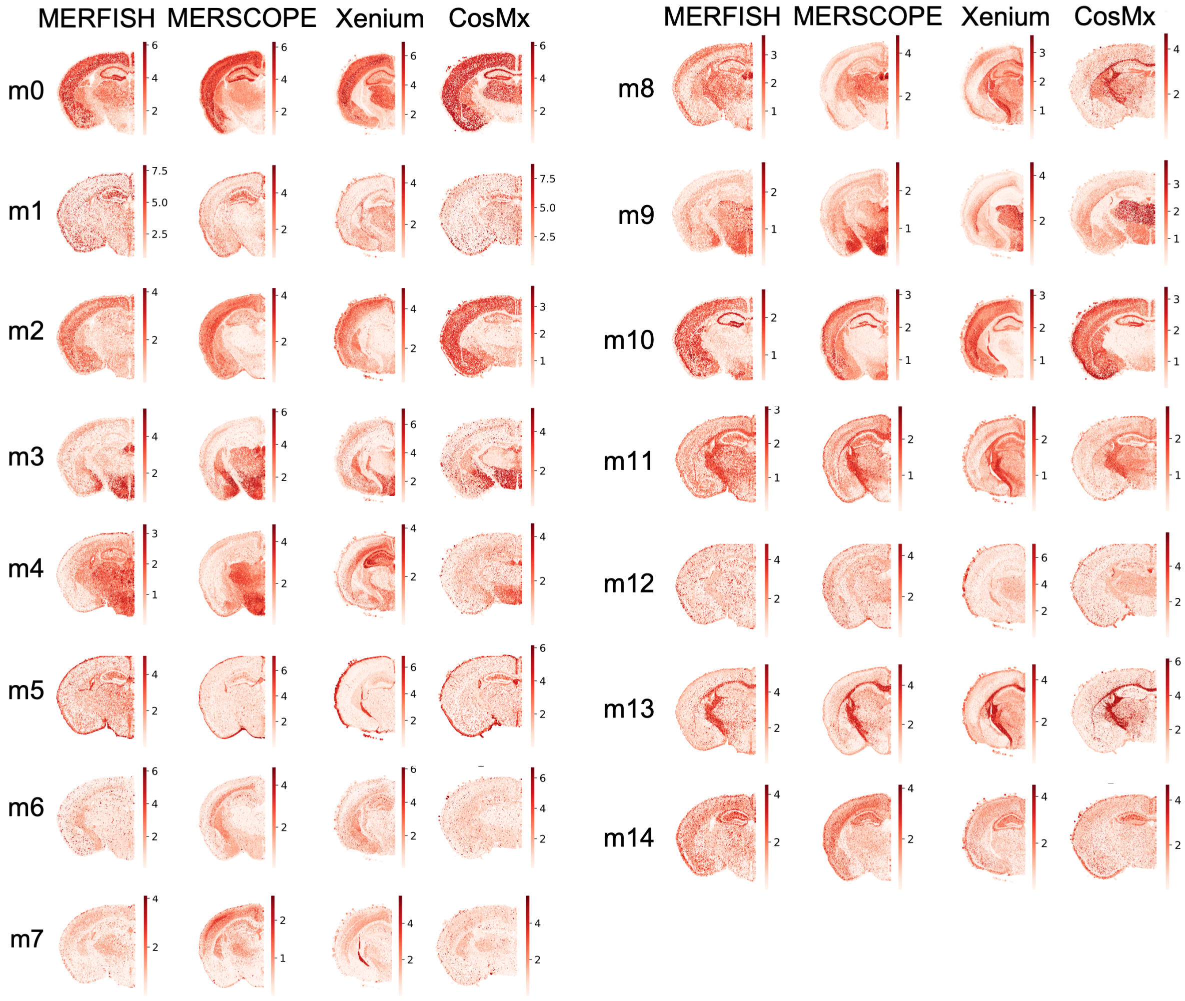


**Figure S2:** *In situ* expression patterns of the 15 metagenes identified by SpiceMix+ Lloki across four spatial transcriptomics technologies: MERFISH, MERSCOPE, Xenium, and CosMx. Each row corresponds to one of the 15 metagenes (m0-m14), and each column displays its spatial expression pattern within a given technology. Darker colors indicate higher expression levels.

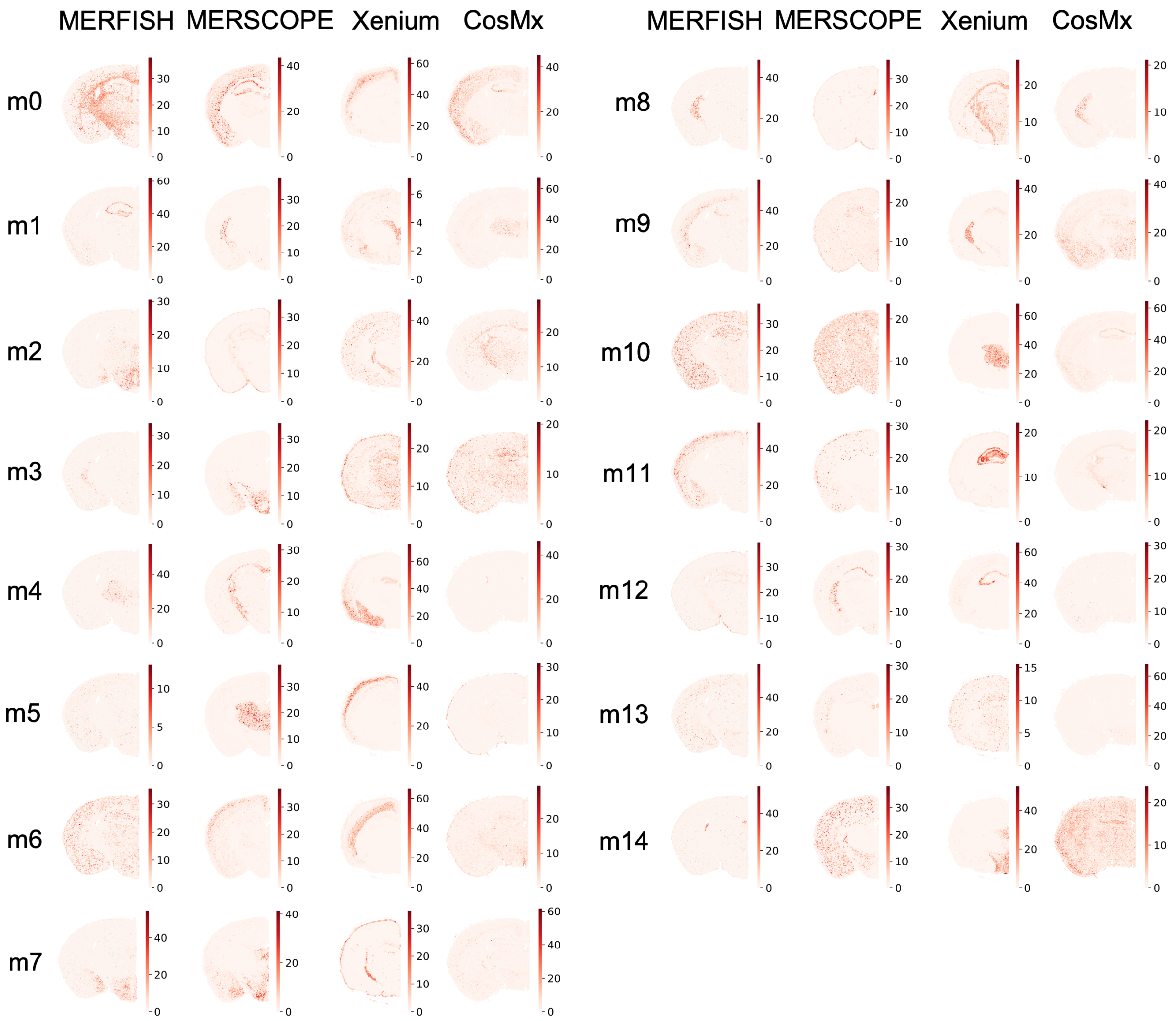


**Figure S3:** *In situ* expression patterns of the 15 metagenes identified by SpiceMix on the shared gene set (22 genes) across four spatial transcriptomics technologies: MERFISH, MERSCOPE, Xenium, and CosMx. Each row corresponds to one of the 15 metagenes (m0-m14), and each column displays its spatial expression pattern within a given technology. Darker colors indicate higher expression levels.

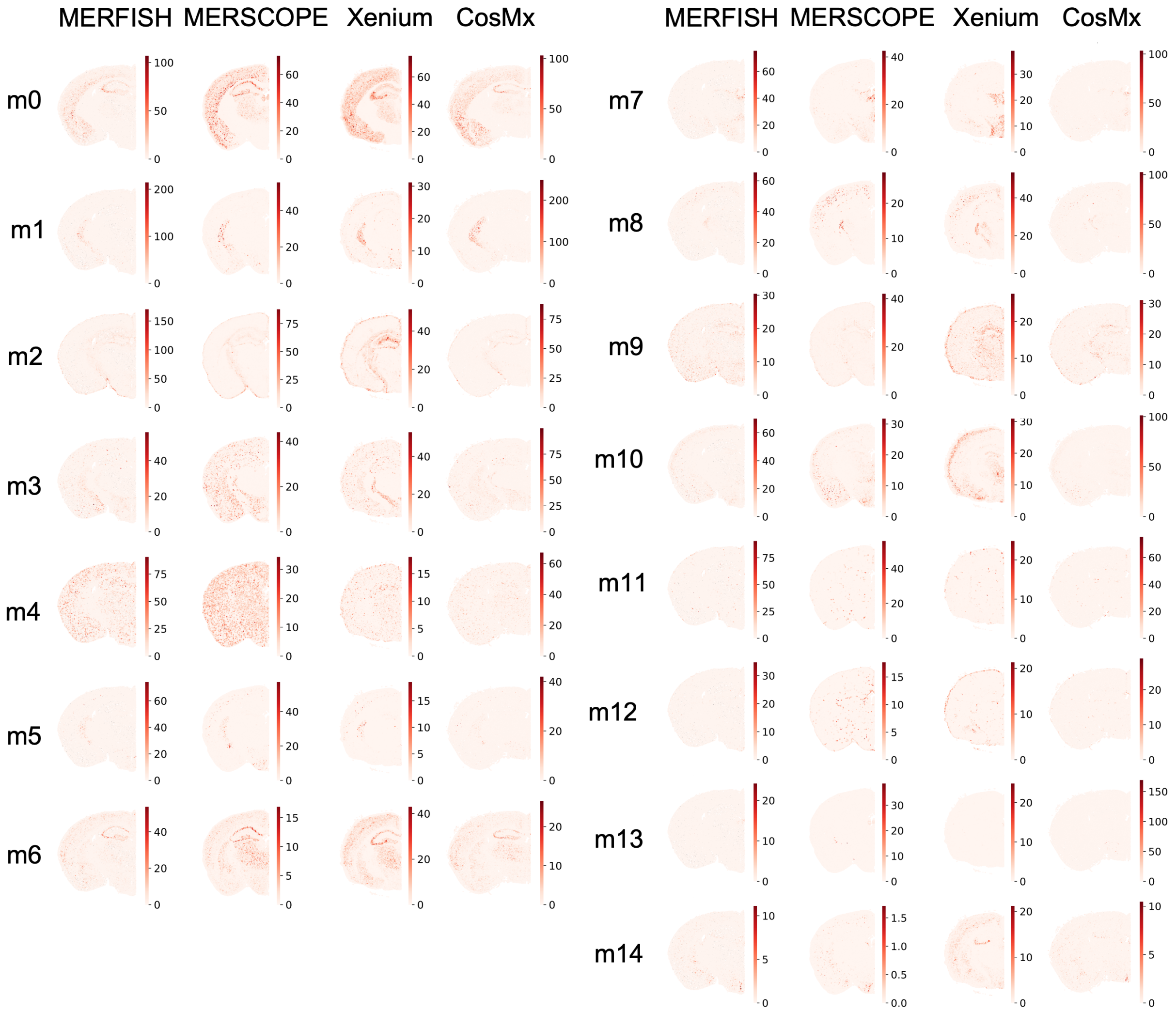


**Figure S4:** *In situ* expression patterns of the 15 metagenes identified by independent SpiceMix runs performed separately on each spatial transcriptomics technology: MERFISH, MERSCOPE, Xenium, and CosMx. Each row corresponds to one of the 15 metagenes (m0–m14), and each column displays its spatial expression pattern of that metagene in the respective technology. Darker colors indicate higher expression levels.
